# Neural Network-Enhanced Investigation of Ferroptosis and Druggability in Early-Onset Alzheimer’s Disease

**DOI:** 10.1101/2025.06.21.660896

**Authors:** Pratibha Singh, Soumya Lipsa Rath

## Abstract

**Background:** Alzheimer’s disease (AD) is a complex neurodegenerative disorder which is multifactorial in nature. Some of its characteristics are slow cognitive decline, memory problems and behavioral changes. AD patient brains show a progressive synaptic toxicity, autophagy, neuroinflammation, excess generation of reactive oxygen species (ROS), neuronal death and oxidative stress, which occurs due to disrupted metal homeostasis along with tau and amyloid-β protein deposition. Notably, lipid peroxidation, iron buildup and elevated oxidative stress in AD brains suggest a possible molecular connection between ferroptosis and AD neurodegeneration.

**Methods:** This study explores the genetic and bioinformatics perspective on the relationship between ferroptosis and AD aiming to identify potential therapeutic biomarkers using Neural network (NN) and Machine learning models. Six ferroptosis related genes were found to be differentially expressed in AD. Further machine learning analysis shortlisted four key biomarker genes. An NN-based diagnostic prediction model was developed and validated using AUC-ROC anaysis, which gave high diagnostic values (AUC-0.92) in the analysis.

**Results:** The findings highlight a strong correlation between ferroptosis and altered metabolic functions in AD. miRNA-gene interaction analysis revealed that two biomarker genes, CYBB and ACSL4 can be regulated by several regulatory miRNAs i.e., hsa-miR-146-5p, hsa-miR-106b-5p, hsa-miR-223-3p, hsa-miR-155-5p, hsa-miR-34a-5p, hsa-miR-125b-5p and hsa-miR-27a-3p suggesting their potential as early diagnostic biomarkers. Immune microenvironment analysis revealed strong neuroinflammatory responses were in AD with increased infiltration of macrophages (M0, M1 and M2), monocytes and multiple T cell subsets. This heightened immune activity may be driven by ferroptosis-induced oxidative stress, contributing to neuronal death. Furthermore, druggability of these targets was evaluated and several drugs were identified that may be potentially repurposed for therapeutic intervention in AD pathogenesis.

**Conclusion:** This study presents a diagnostic predictive model integrating gene expression, miRNA regulation and immune infiltration analysis, offering a novel perspective on early AD detection. The identified ferroptosis-related biomarkers and regulatory miRNAs could serve as valuable tools for clinical diagnosis and targeted therapeutic intervention, advancing personalized treatment strategies for Alzheimer’s disease.

## Introduction

Alzheimer’s disease (AD) is the most common form of dementia worldwide, presenting a major challenge to healthcare systems and society. AD is a multifactorial complex neurodegenerative disorder, characterized by slow cognitive degradation, memory malfunction, and behavioral transformations. AD patients’ brains show progressive synaptic toxicity, autophagy, neuroinflammation, excess generation of reactive oxygen species (ROS), neuronal death, and oxidative stress due to an imbalance in metal homeostasis, in addition to the deposition of tau and amyloid-β protein [1, 2]. Critical brain functions like myelin formation, neuronal activity, neurotransmitter production, and energy metabolism are impacted by iron, which is a transition metal [3, 52]. According to studies, an imbalance in iron homeostasis might prevent the mitochondrial electron transport chain from functioning, which results in oxidative stress [4]. Increased oxidative stress, lipid peroxidation, and iron buildup in AD brains points to a possible molecular connection between ferroptosis and AD neurodegeneration [5]. Ferroptosis is defined as a type of programmed cell death mediated by iron-dependent lipid peroxidation that is different from autophagy, apoptosis, and necrosis [6]. It is specifically characterized by an iron-dependent accumulation of lipid peroxides [7]. Ferroptosis is primarily dependent on two physiological processes: degradation (particularly autophagy and the ubiquitin-proteasome system) and cell metabolism (particularly lipids, iron and amino acids) [8].

For many years, traditional diagnostic strategies have encompassed clinical examination, cognitive testing, neuroimaging, and biomarker analyses to detect AD [9]. These methods, however, come with their own limitations, ranging from subjectivity to relatively low sensitivity and invasiveness [10]. Current diagnostic models include identifying people affected by AD by clinical suspicion based on cognitive assessment, neuropsychological testing, and then by neuroimaging (such as MRI, PET imaging) and biofluid assessment (such as cerebrospinal fluid analysis) [11]. Although these methods allow certain useful characterizations of AD pathology, there are certain limitations [12]. Adherence to protocols for clinical assessments and cognitive tests is subjective and predisposes these assessments to inter-rater variability, resulting in inconsistent diagnosis [13]. Additionally, some of the neuroimaging techniques may be insensitive to early-stage AD changes. Although neuroimaging techniques like neuromelanin-MRI of the locus coeruleus can provide early-stage AD detection but are limited by focusing only on norepinephrine and neglecting other key neurotransmitters like serotonin and acetylcholine which may have potential roles in neuropsychiatric symptoms (NPS) system [14]. Several invasive modalities of biomarker analysis, such as biopsy, can be too risky and not feasible, nor always universally accessible to all patients [15]. Over the past few years, ML and neural network-based approaches have shown great potential for a paradigm shift in AD diagnostics [16]. Importantly, the use of these novel computational tools promises improved accuracy, efficiency, and scalability to drive new discoveries in the fight against AD [17].

AD is a degenerative neurological ailment that worsens over time and for which there is presently no recognized cure. It is essential to create efficient algorithms for illness diagnosis. In some cases, data-driven diagnostic and predictive approaches for AD are accomplished through the application of machine learning (ML) techniques [18]. Research utilizing ML approaches has demonstrated impressive accuracy rates in distinguishing between patients with dementia and healthy controls, as well as between various forms of dementia [19]. However, more investigation is required to determine the therapeutic usefulness of machine learning in dementia diagnosis. This includes assessing the cost-effectiveness and potential effects of these approaches on patient outcomes [20].

Recent research on AD has explored various approaches to understand its characteristics, causes, and potential treatments, including the use of artificial intelligence (AI) for early detection [21]. Studies have examined AI applications such as speech analysis and voice biomarkers for dementia detection [22]. However, the findings have been limited by variations in research populations and data collection methods, highlighting the need for standardized procedures [23]. Multimodal approaches that combine different data types, such as imaging and non-imaging modalities, have been proposed to enhance the accuracy of early AD diagnosis, but the lack of standardized data collection and analysis methods poses a major drawback [24]. Cognitive evaluation tools, including the mini-mental state examination and neuroimaging techniques, have been utilized for early detection of AD [25]. This also suggests that integrating cognitive assessments with early diagnostic methods could be effective [26]. However, these tools often lack the sensitivity required for precise detection, necessitating the development of more accurate instruments [27]. ML algorithms have shown high accuracy and sensitivity in distinguishing AD patients from control groups, but most often these studies are limited by small sample sizes, requiring validation in larger and more diverse populations [28]. While AI’s potential in healthcare, particularly in cognitive problem diagnosis, is recognized, there is a need for more in-depth exploration of specific methodologies and consideration of the costs and financial implications in implementing these technologies [29]. Additionally, research using ensemble ML and bioinformatics has identified novel biomarkers and pathways for AD, highlighting the importance of these in-silico methods while acknowledging challenges related to inconsistencies in study methods, demographic factors and data collection methods [30]. There are a number of unexplored areas in the field of AI-based AD diagnosis. To increase diagnostic accuracy, integrating several data types such as MRI, PET, EEG/MEG and sensor data is crucial [31]. Many of the current research studies concentrate on classifying AD using various ML methods [32]. Moreover, some of them also provide early diagnosis of AD but with certain limitations [33]. Our study addresses the existing limitations and problems in the field of AI-based approaches for cognitive decline detection by offering the early diagnosis of AD utilizing ML and NN-based methodologies [34].

In this study, we will present the genetic and bioinformatics perspective on the relationship between ferroptosis and AD using neural network and machine learning models. Our objective is to target ferroptosis-related genes that were abnormally expressed in AD to uncover potential miRNAs which can be used for further diagnosis and treatment [35]. In order to evaluate the hub gene’s ability to detect AD and their association with immune infiltration in AD patients, these genes were used to build diagnostic models [36]. Finally, we can offer important data support for the clinical treatment and therapeutic development of AD by screening microRNAs and hub genes [37].

## Materials and Methods

### 1. DATA SET COLLECTION

The Gene Expression Omnibus (GEO) database was used to download the AD dataset GSE118553, and microarray dataset GSE33000 and GSE157239 on AD were retrieved for confirmatory research [38]. Using the FerrDB [39], Kyoto Encyclopedia of Genes and Genomes (KEGG) [40], Gene Set Enrichment Analysis (GSEA) [41], and relevant literature, 106 genes associated to ferroptosis were identified.

### 2. DIFFERENTIAL EXPRESSION ANALYSIS USING Z-SCORE NORMALIZATION AND FDR CORRECTION

Raw gene expression data from the GSE118553 dataset were retrieved from the NCBI GEO using the GEOquery package in R [42]. Expression matrices were extracted and preprocessed to ensure consistent gene annotation and sample alignment. To standardize expression profiles across samples, z-score normalization was applied gene-wise using the formula [45]:

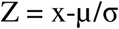

Where, x is the expression value for a gene, μ is the mean expression across all samples and σ is the standard deviation. Genes with absolute z-scores greater than 1.96 were considered statistical outliers, corresponding to a 95% confidence level under the standard normal distribution.

To identify significantly dysregulated genes, nominal p-values obtained from differential expression analysis were adjusted for multiple testing using the Benjamini–Hochberg False Discovery Rate (FDR) method [49]. Genes meeting both criteria |z| > 1.96 and FDR-adjusted p-value < 0.05 were considered significantly differentially expressed. (z > +1.96) and downregulated (z < –1.96) genes were quantified and z-score distributions were visualized to assess overall gene expression variability.

To ensure the robustness of the differentially expressed genes (DEGs) identified through z-score normalization, a parallel analysis was performed using the Linear models for microarray data (Limma) (version 3.60.4) package in R. This linear modeling approach provided an independent statistical framework to cross-validate DEG detection with A p-value of less than 0.05 and |log2 fold change (FC)|>0.5 was taken as the relaxed criteria in this linear modeling approach to provide an independent statistical framework to cross-validate DEG detection. This aligns with accepted practices in transcriptomic analysis, where moderate fold changes may still reflect meaningful biological signals and can be functionally significant [50].

### 3. PATHWAY ENRICHMENT ANALYSIS

KEGG Pathway gene annotations were obtained via the KEGG rest API to conduct GSEA [41]. Genes were plotted against the background set using the GO annotations found in the R-package3 (v.3.1.0) [43]. The clusterProfiler (v 3.14.3) R-package was utilized for enrichment analysis to derive the GSEA results [44]. On the basis of phenotypic grouping and expression profiles minimal gene set was maintained at two and the maximum gene set at five thousand respectively. Phenotypic grouping involves categorizing samples on the basis of phenotypes i.e., observable traits or characteristics. Here, phenotypic groups include individuals with early stage AD, advanced stage AD or no AD (control samples). On the other hand, levels at which genes are expressed in the given sample is given by Expression profiles, which helps in identifying the and downregulated genes in specific phenotypes. Using minimal and maximal limits to gene sets ensures inclusion of only relevant pathways eliminating all other insignificant pathways as well as non-specific gene sets which could dilute the relevance of results. Statistically significant result was defined as those with a p-value of less than 0.05 and an False Discovery Rate (FDR) of less than 0.1. To investigate the biological roles and associated pathways of DEGs, GO and KEGG pathway enrichment analysis were carried out on DEGs [45].

### 4. CONSTRUCTION OF CLASSIFIER MODEL

Data on gene expression alongwith the time and status of survival were integrated using the R-package, glmnet and RandomForest [46]. Regression analysis was done using the lasso-cox (Least Absolute Shrinkage and Selection Operator) [47] and Random Forest (RF) techniques. These were used to perform feature selection for the construction of a diagnostic model using the most representative genes. To find the best model, 10-fold cross-validation was additionally set up. The outcomes of the two types of machine learning models namely Lasso and RF were cross-screened to develop the final diagnostic prediction model.

### 5. VALIDATION OF DIAGNOSTIC MODEL

Receiver Operating Characteristic (ROC) analysis was performed using pROC in the R-package to produce the Area Under the Curve (AUC) [48]. In order to get the final AUC values, AUC and confidence intervals (CIs) were also analyzed using the pROC CI function. The entire process assisted in observing the expression of distinctive genes in the AD dataset GSE118553.

### 6. SUBGROUP ANALYSIS

To complete the cluster analysis, ConsensusClusterPlus was used. Here agglomerative pam clustering with a 1-spearman correlation distance and resampling 80% of the samples for ten repeats were used to carry out the analysis. Unsupervised hierarchical clustering was carried out using the gene expression of potential genes as input and R’s “ConsensusClusterPlus” [51]. Subgroup DEGs were obtained by applying Limma analysis to the various subgroups, and KEGG and GO were utilized to assess the functional differences among the subgroups. Networkanalyst4 was used to build a gene-miRNA interaction network in order to predict the miRNAs of potential genes [52].

### 7. ANALYSIS OF IMMUNE-MICROENVIRONMENT

We used the CIBERSORT method, based on our gene expression profiles, to estimate the scores of 22 different immune-infiltrating cell types for each sample [53]. This analysis was performed using the Immuno-oncology biological research (IOBR) R package [54]. To assess immune cell infiltration, we employed CIBERSORT within the R environment, calculated correlations using the Spearman coefficient, and visualized the relationships between infiltrating immune cells through a heat map generated with the corrplot package [53].

### 2.8. HUB GENES-miRNA NETWORK CONSTRUCTION

Networkanalyst4 was used to build a gene-miRNA interaction network in order to predict the miRNAs of potential genes [55]. Here, genes and hub miRNAs are identified as potential biomarkers for therapeutic purposes.

## Results

### 1. Pathway enrichment analysis of differentially expressed genes (DEGs) shows predominance of Ferroptosis pathway in up regulated genes

The GEO database was used to download the AD dataset GSE118553. This database is based on analysis of the transcriptome in likely asymptomatic and symptomatic Alzheimer brains. The human brains of 27 control, 33 Asymptomatic AD (AsymAD), and 52 AD participants were microarray profiled, and both differential and co-expression analyses were carried out. AsymAD refers to those who have intact cognition and neuropathology suggestive of AD. These people have a high likelihood of developing AD, however it is still uncertain whether transcriptome changes in the brain could reveal mechanisms underlying their vulnerability to AD. Cerebellum is one of the tissues that is largely unaffected by the disease, however frontal, temporal, and entorhinal cortical regions are known to be affected by AD neuropathology. The frontal cortical area in AD patients may be the starting point for basic brain alterations, according to transcriptome activity in AsymAD participants. Furthermore, this dataset provides new insight into the initial biochemical changes occurring in the brain prior to clinical AD diagnosis, opening up new pathways for therapeutic interventions aimed at preventing AD [56].

To identify genes with significant expression alterations in the GSE118553 dataset, z-score normalization was applied across the entire gene expression matrix. This approach standardized gene expression values, enabling the identification of statistical outliers using the threshold of |z| > 1.96, corresponding to the 95% confidence interval of a standard normal distribution. From a total of 18,976,523 gene sample expression values, 8,73,873 values (4.61%) were identified as expression outliers. Among these, 47,323 values exhibited significant upregulation (z > +1.96), indicating elevated expression relative to the global mean. To further refine the selection of biologically significant genes, raw p-values were corrected for multiple testing using the Benjamini–Hochberg False Discovery Rate (FDR) procedure. Genes satisfying both the FDR-adjusted p-value < 0.05 and |z| > 1.96 criteria were classified as significantly differentially expressed. This dual-filtering strategy improved the specificity of detection by incorporating both expression magnitude and statistical confidence. The distribution of z-scores across all genes showed a near-normal curve with prominent tails representing potential biologically relevant dysregulation. These results highlight a subset of genes with significant transcriptional deviations that may contribute to the underlying biological differences captured in the dataset (Fig. 1(A)). We also identified a total of 3,750 DEGs in the dataset corresponding to AD utilizing the Limma method while cross-validating the DEGs obtained from z-score followed by FDR correction, out of which 2,107 genes were up-regulated and 1,643 were downregulated (SI. 1). In differential expression of genes, some genes show increased while others show decreased expression levels under specific conditions or in response to certain stimuli. When compared to a control group, these genes are termed as up-regulated and downregulated genes respectively. The up-regulated genes are crucial for many biological functions and can reveal important information about how cells react to stimuli and how diseases are caused, hence only up-regulated genes were considered for further processing. The candidate genes were found to be predominantly enriched in the pathways for “Ferroptosis,” “Fatty acid degradation,” and “Focal adhesion” (Fig. 1(B)). This was identified by using KEGG pathway analysis. On the basis of phenotypic grouping and expression profiles, we maintained the minimal gene set at five and the maximum gene set at five thousand during the pathway analysis. Among the three obtained pathways, Ferroptosis pathway covered a relatively larger proportion among the candidate genes.

**Fig. 1.**
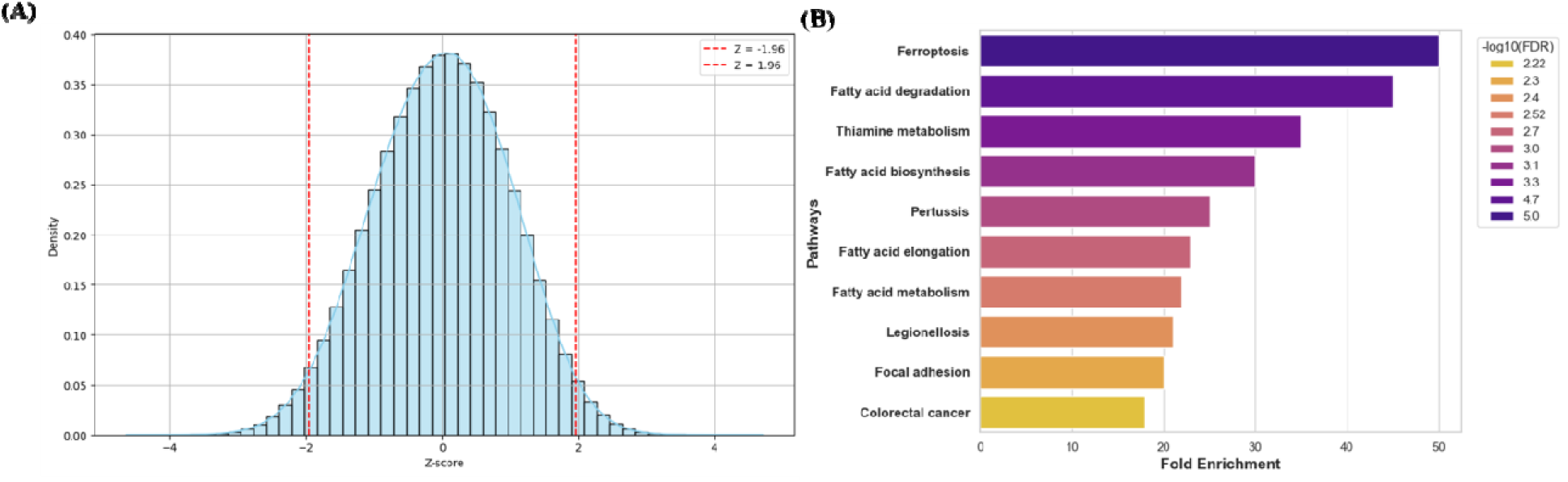
**(A)** Distribution of z-score normalized gene expression values across the Alzheimer’s disease dataset (GSE118553). Vertical red dashed lines indicate ±1.96 thresholds used to define significant upregulation and downregulation, **(B)** KEGG analysis pathways associated with DEGs obtained from Alzheimer’s dataset (*x-axis: Fold Enrichment; y-axis: Enrichment pathways)*.

According to Gene Ontology (GO) analysis, candidate genes were primarily found in “mitochondria”, “organelle envelope” and “envelope” in terms of cellular components (Fig. 2(A)). The “positive regulation of multicellular organismal process”, “cell motility”, “localization of cells” and other transport-related processes make up the main biological functions of the candidate genes (Fig. 2(B)). The three most significant roles of the candidate genes, as determined by molecular function (MF), were “oxidoreductase activity”, “heme binding” and “iron ion binding” all of which are closely associated with ferroptosis (Fig. 2(C)). From the GO and KEGG pathway analyses, the insights on different biological roles and other pathways associated with the DEGs were obtained from AD dataset. Results indicate that there is a strong molecular connection between Alzheimer’s disease and ferroptosis process.

**Fig. 2.**
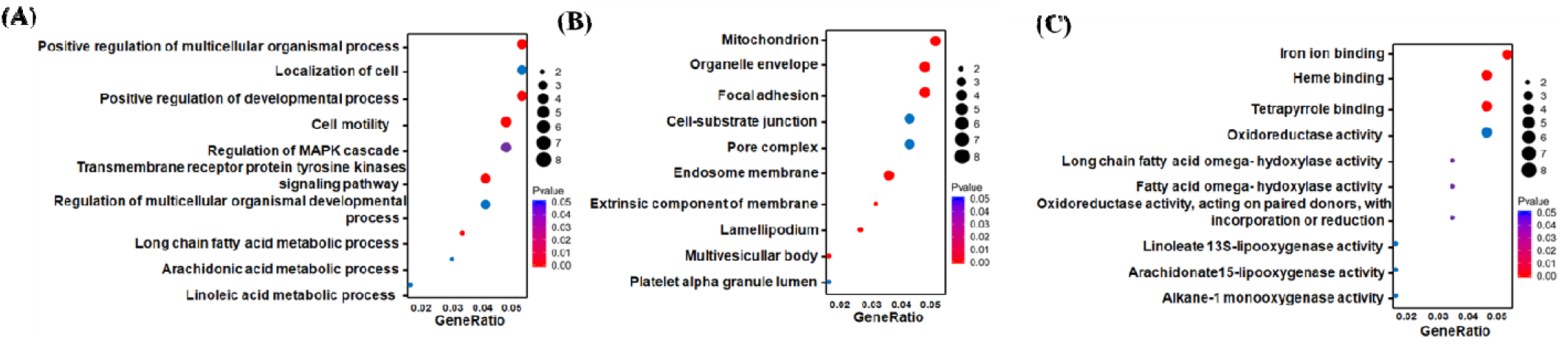
**(A)** GO analysis of potential genes for cellular distribution *(x-axis: GeneRatio; y-axis: Cellular distribution),* **(B)** GO analysis of potential genes for biological processes *(x-axis: GeneRatio; y-axis: Biological processes)* and **(C)** GO analysis of potential genes for molecular function *(x-axis: GeneRatio; y-axis: Molecular Functions). Here, redder dots indicate more highly enriched terms (lower FDR/p-values, stronger confidence) and bluer/purpler dots indicate borderline significant or less confident terms*.

### 2. Identification of candidate genes using Machine learning models

#### 2.1. Potential candidate genes were identified utilizing supervised ML models on AD dataset and ferroptosis related genes

In the previous section, up-regulated differentially expressed genes alongwith their associated pathways and functions were identified. In this section, we have also included genes that are linked to ferroptosis. 106 genes associated to ferroptosis were identified using the FerrDB, KEGG, GSEA, and relevant literature. Subsequently, a set of common genes were obtained after performing a crossover on both the gene sets of AD DEG and ferroptosis related genes (SI. 2). According to previous comparative studies done on performance and accuracy of various ML models (like Lasso, RF, Support Vector Machine (SVM) and Decision Tree), Lasso and Random Forest have been found to give exceptionally good models with high accuracy [46, 47]. Hence, two ML models viz., Lasso and RF were developed using the obtained common genes. Data on gene expression alongwith the time and status of survival were integrated using the R-package, glmnet and RandomForest. In order to identify potential genes, LASSO regression was first utilized. LASSO is a supervised regularization technique. The AD dataset yielded the identification and determination of six putative candidate genes namely CYBB (Cytochrome b-245 beta chain), AKR1C1 (Aldo–Keto Reductase 1), ACSL4 (Acyl-CoA synthetase long-chain family member 4), GPX4 (Glutathione Peroxidase 4), FTH1 (Ferritin Heavy Chain 1) and MAPT (Microtubule Associated Protein Tau) (Fig. 3(A)). Additionally, we employed RF regression which is also a supervised ML technique to find candidate genes, from which four possible candidate biomarkers were identified namely CYBB, ACSL4, GPX4 and AKR1C1 (Fig. 3(B)). Both the models were then used to perform feature selection for the construction of a diagnostic model using the most representative genes. To find the best model, 10-fold cross-validation was additionally set up. After cross-analyzing the genes found by these two ML algorithms, four potential genes (CYBB, ACSL4, GPX4 and AKR1C1) were identified which can be used to develop the final diagnostic prediction model. We verified the diagnostic value of these four candidate genes using ROC curves at the point in the AD dataset. All candidate genes were employed as joint indicators and the obtained values were (AUC 0.92, CI 0.95) thus, indicating a very high diagnostic significance (Fig. 3(C)). Furthermore, an investigation of these potential biomarker gene’s expression profiles was also carried out. The violin plot in Fig. 3(D) shows the expression distribution of four genes (CYBB, ACSL4, GPX4 and AKR1C1) across AD (yellow) compared to healthy controls (orange). The expression levels are represented on the y-axis with separate violin plots for each condition i.e., AD and healthy alongwith -log (p) values as statistical significance for each pair which is shown in the form of asterisk (*). Distribution shapes and central tendency markers indicates that CYBB and ACSL4 exhibit noticeable increase in expression in the AD group compared to controls which is consistent with statistically significant differences (p < 0.05). In contrast, GPX4 and AKR1C1 shows relatively similar expression or little variation in distributions between the two groups indicating no significant change. This suggests that they did not undergo meaningful transcriptional alterations in AD while CYBB and ACSL4 are differentially expressed. These pronounced differences in expression are suggestive of involvement of these genes in pathological processes underlying AD which could further serve as potential biomarkers or therapeutic targets. The entire process assisted in observing the expression of distinctive genes in the AD dataset GSE118553.

**Fig. 3.**
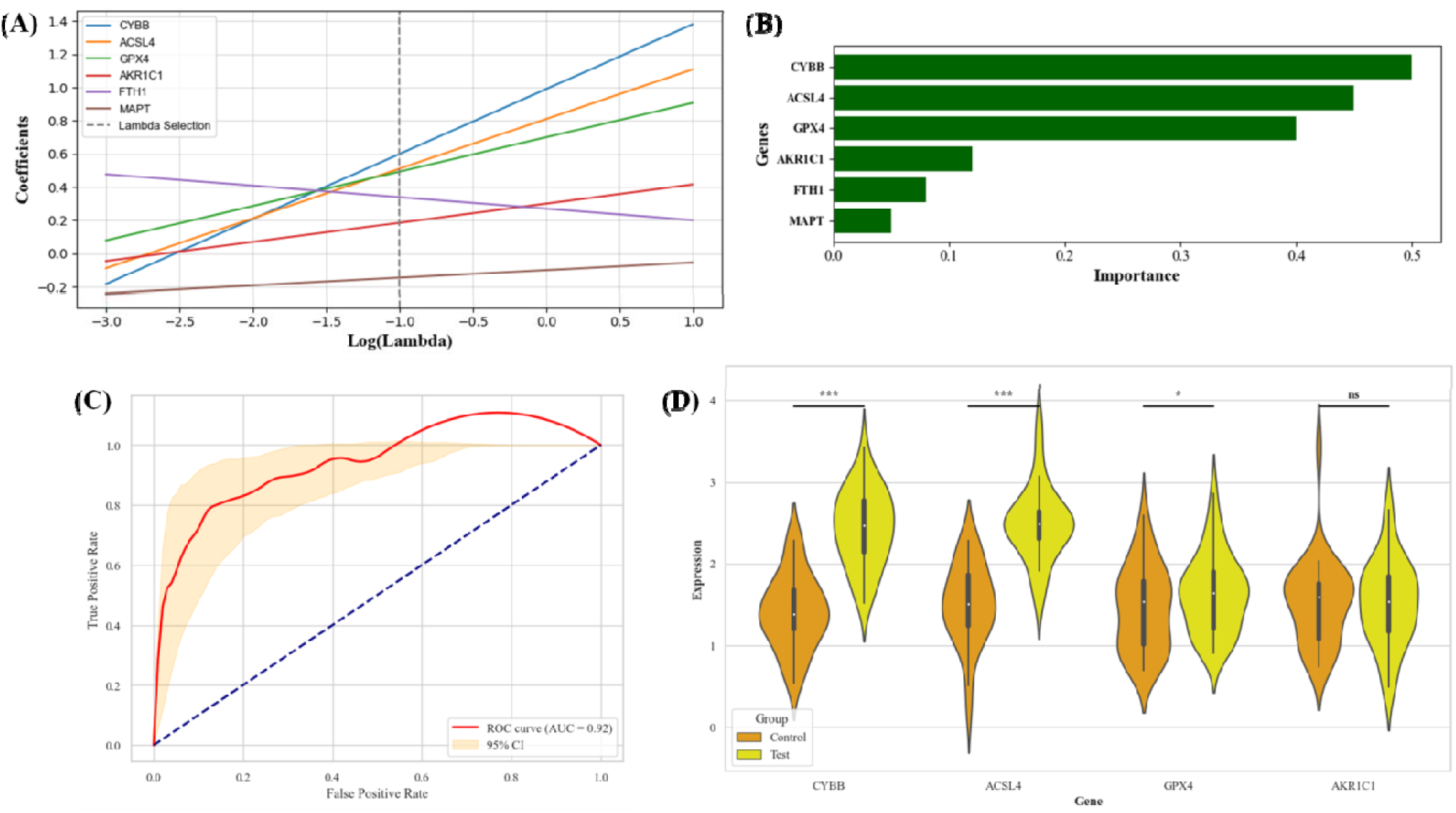
**(A)** Candidate gene identification for Alzheimer’s disease dataset using Lasso regression, **(B)** Candidate gene identification for Alzheimer’s disease dataset using Random Forest (RF) regression curve, **(C)** validation using ROC curve for Alzheimer’s dataset, **(D)** Analysis of candidate gene expression profile for AD dataset using violin plot with asterisks indicating statistical significance estimated based on expression trends*: * p < 0.05 (borderline), ** p < 0.01, *** p < 0.001, ns = not significant*.

#### 2.2. Unsupervised ML models were used to confirm the association of relevant candidate genes with both AD and ferroptosis

We conducted consensus clustering (CC) analysis on the AD dataset GSE118553, taking intra-group consistency into consideration. By averaging the intra-group consistency of clusters, we obtain the maximum number of clusters, i.e., K = 2. Based on the clustering chart, we discovered that the clustering between groups was strongest at K = 2, which led to the formation of the two unsupervised clustering subgroups, C1 and C2 (SI. 3(A), (B)). Violin plots were used to show the candidate gene expression levels in the two clusters (Fig. 4(B)). The expression distributions for GPX4 and AKR1C1 appear similar between the two clusters (with p-values of 0.25 and 0.28 respectively), indicating no statistically significant difference in expression levels between the groups. In contrast, CYBB and ACSL4 shows a prominant difference in expression with Cluster C2 exhibiting higher expression levels than Cluster C1. This difference is statistically significant, supported by a p-value of 0.001. Overall, these results suggest that as observed earlier, here also the expression varies significantly for CYBB and ACSL4 and showed considerable fluctuations between the two clusters, while GPX4 and AKR1C1 remain relatively unchanged.

**Fig. 4.**
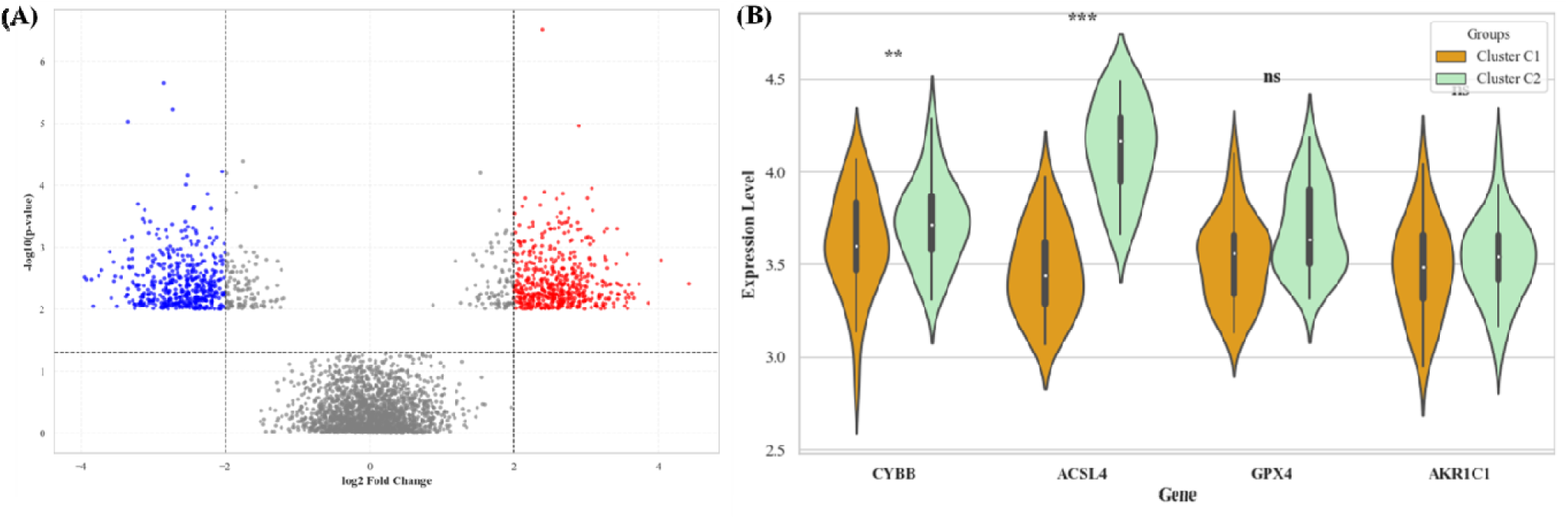
**(A)** Volcano map showing DEGs obtained from subgroup analysis, **(B)** Analysis of differential expression of relevant candidate genes between subgroups using violin plot with asterisks indicating statistical significance estimated based on expression trends*: * p < 0.05 (borderline), ** p < 0.01, *** p < 0.001, ns = not significant*.

After performing limma analysis on the two subgroups, 827 DEGs in total were found. 523 were up-regulated (shown in red color) and 304 were a down-regulated genes (Shown in blue color) (Fig. 4(A)). This shows the efficiency of the proposed diagnostic predictive model.

According to GO analysis, the DEGs were mostly found in the “focal adhesion” and “cell substrate junction” regions of the cell (Fig. 5(A)). The “long chain fatty acid metabolic process” and “arachidonic acid metabolism” are the main biological processes linked to DEGs (Fig. 5(B)). The most important DEGs, according to MF analysis, were “heme binding” and “iron ion binding” (Fig. 5(C)). The functional analysis of the subgroups obtained from unsupervised learning confirmed the association of relevant candidate genes with both AD and ferroptosis.

**Fig. 5.**
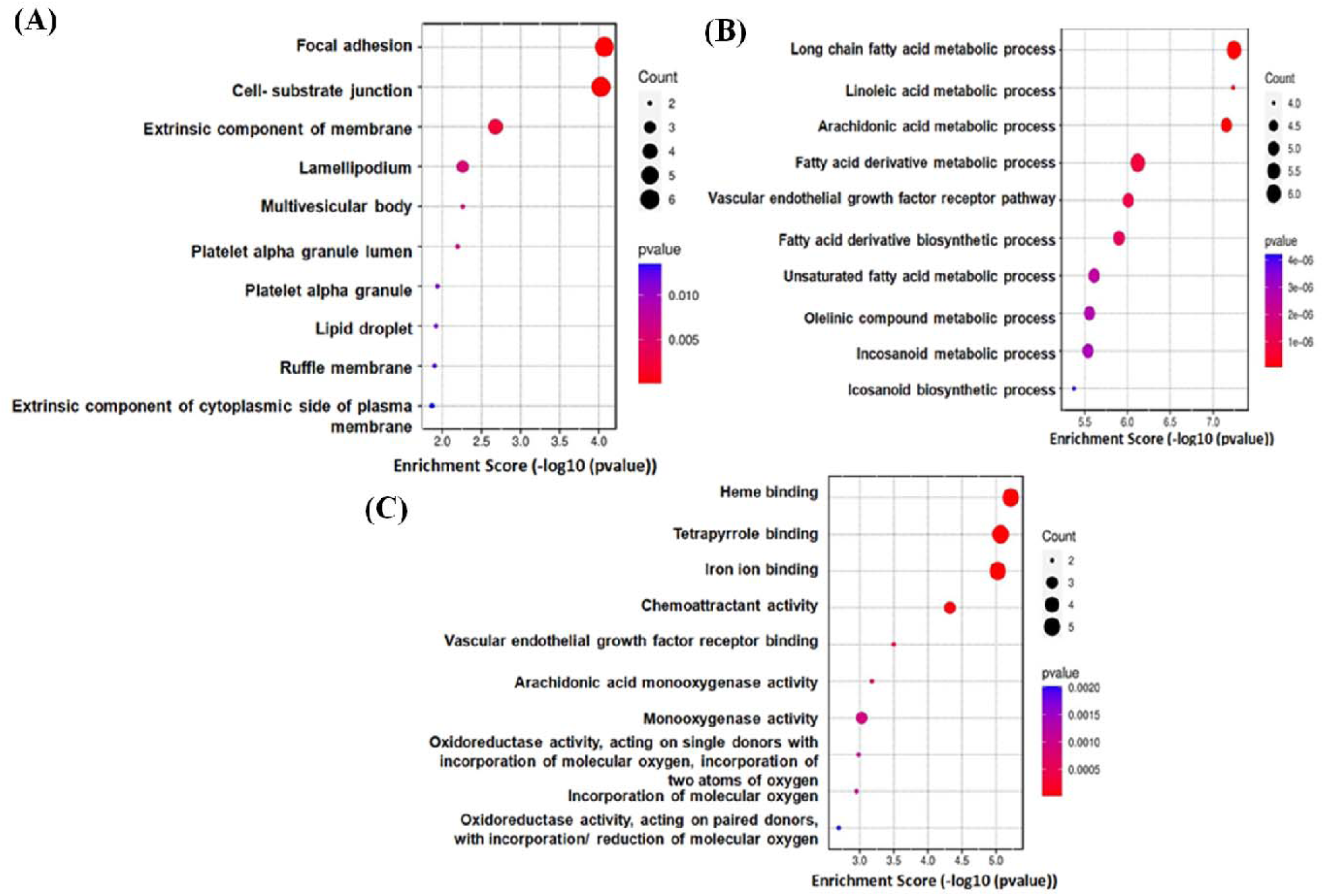
**(A)** Candidate GO analysis of candidate genes for cellular composition and distribution, **(B)** GO analysis of candidate genes for biological processes, **(C)** GO analysis of candidate genes for molecular function. *Here, redder dots indicate more highly enriched terms (lower FDR/p-values, stronger confidence) and bluer/purpler dots indicate borderline significant or less confident terms*.

### 3. Controlling expressions of these candidate genes is an improved therapeutic strategy for early onset of Alzheimer’s disease (AD)

Using Networkanalyst, we created miRNA-gene interaction networks. A gene-miRNA network visually represents the interactions between genes and associated numerous miRNAs. miRNAs can regulate several genes, making these networks useful resources for biological research. Based on their centrality or significance within the network, the network can assist in prioritizing miRNAs or genes to concentrate on for more validation. Moreover, by examining gene-miRNA networks, disease-associated dysregulated miRNAs and their target genes can be identified, offering valuable information on the underlying molecular mechanisms of the disease and suggestive of a potential therapeutic strategy by controlling their expressions. It may also be used as a biomarker for prognosis, diagnosis, or response to treatment [54]. Fig. 6(A) illustrates a directed gene–miRNA interaction network between two AD-associated genes (CYBB and ACSL4) and their shared regulatory miRNAs (hsa-miR-146-5p, hsa-miR-106b-5p, hsa-miR-223-3p, hsa-miR-155-5p, hsa-miR-34a-5p, hsa-miR-125b-5p and hsa-miR-27a-3p). The network structure reveals that CYBB and ACSL4 are targeted by several common miRNAs including miR-34a-5p and miR-155-5p, which act as central hubs linking both genes. These two miRNAs are highlighted in yellow to indicate their elevated betweenness and degree centrality which suggest their importance in gene regulatory interactions. The bar plot shown in Fig. 6(B) quantifies node centrality, indicating that CYBB and ACSL4 possess the highest centrality scores which in turn reflects their prominent roles as major regulatory targets in this network. Among miRNAs, miR-155-5p and miR-34a-5p exhibit the highest centrality indicating their potential as key post-transcriptional regulators in AD-related pathways. Other miRNAs, such as miR-146a-5p, miR-223-3p, miR-125b-5p, miR-27a-3p and miR-106b-5p shows moderate centrality, indicating secondary but potentially synergistic regulatory roles. Overall, the network analysis reveals a tightly coordinated miRNA-mediated regulatory mechanisms, with certain miRNAs potentially influencing cross-regulation of the oxidative stress and lipid metabolism pathways via CYBB and ACSL4.

**Fig. 6.**
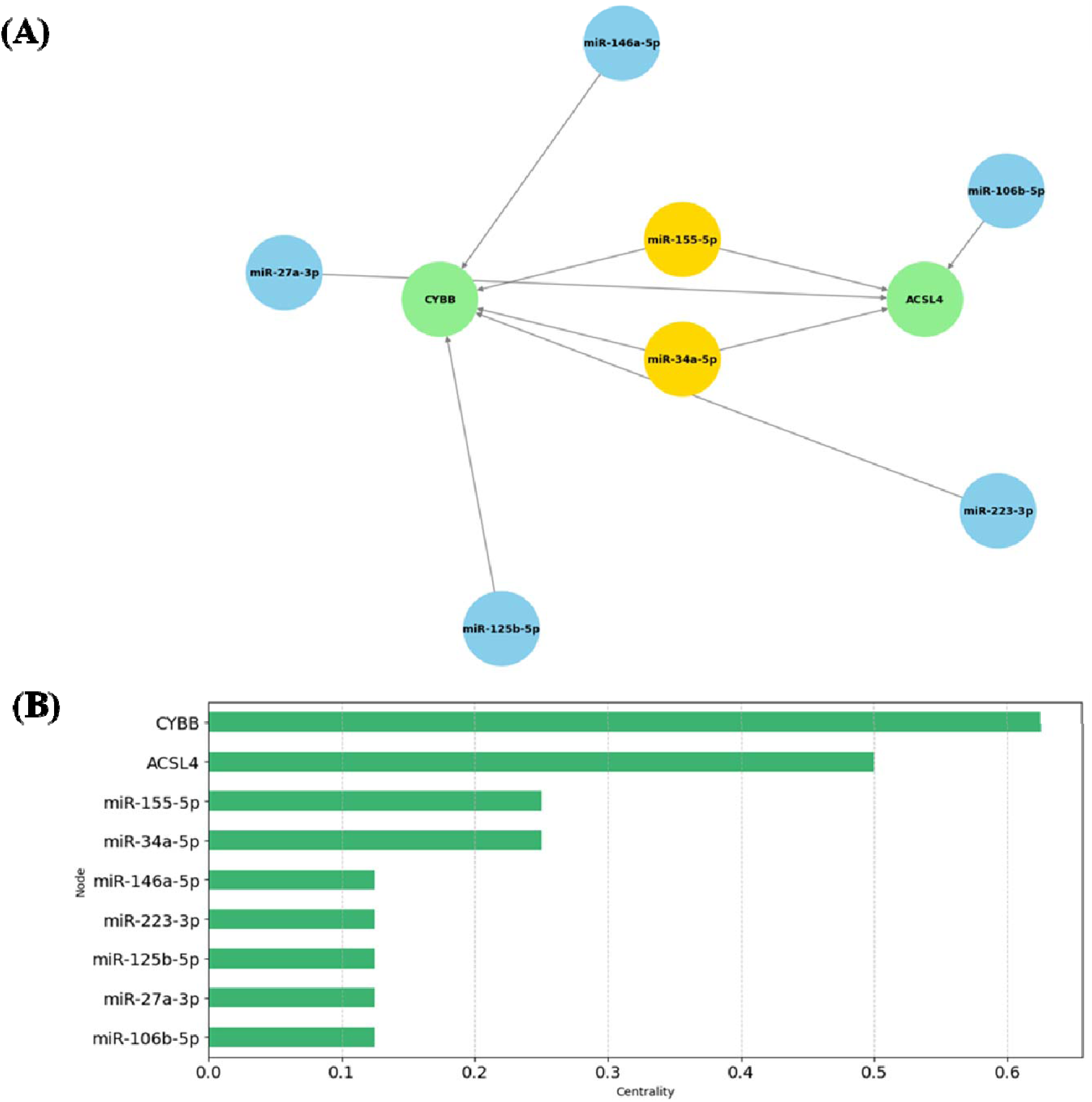
**(A)** Network showing interactions between candidate genes and miRNAs, **(B)** Degree Centrality plot for nodes in the interaction network.

CYBB contributes to the generation of ROS through NADPH oxidase complex, which has been linked to oxidative stress, a major contributing factor to AD [57]. Similarly, ACSL4 is involved in the production of polyunsaturated fatty acids, which is a process related to lipid metabolism. Dysregulation of lipid metabolism is also linked to the etiology of AD [58, 59]. AKR1C1 and GPX4 are not emphasized here because they does not exhibit notable expression changes in the diseased state being studied. Furthermore, the factors or certain pathological circumstances like oxidative stress and lipid peroxidation that cause AD may have a lesser impact on GPX4 and AKR1C1 expression than CYBB and ACSL4 which suffers up and down regulations because of the aforementioned factors.

Additionally, research suggests that GPX4 and AKR1C1 may be associated with a more chronic or later-stage process which is not the primary focus in the current analysis, whereas CYBB and ACSL4 are a part of an acute response to early pathology alterations in Alzheimer’s [60].

One important modulator of disease gene expression is micro RNA or miRNA. hsa-miR-146a-5p is one of the most extensively studied miRNAs in AD, which increases in the brains of people with AD, especially in disease-affected regions like the cortex and hippocampus. It is important for controlling neuroinflammation and the innate immune response, two processes that are fundamental to the pathology of Alzheimer’s disease. Although miR-146a-5p may serve as a defense mechanism to regulate excessive inflammation, its persistent upregulation in Alzheimer’s disease may lead to prolonged neuroinflammatory responses that accelerate neurodegeneration [61]. hsa-miR-155-5p is responsible for modulating the immune responses and oxidative stress and thereby preventing neuroinflammation and neuronal damage [62, 63]. hsa-miR-34a-5p is a critical regulator of neuronal apoptosis and synaptic function and targets several AD-related genes alongwith oxidative stress modulators [64, 65]. hsa-miR-27a-3p regulates both chronic inflammation and dysregulation of lipid metabolism, which are two key elements of AD pathogenesis. It also has a role in regulating synaptic activity, which is important in AD, because synapse loss correlates significantly with cognitive decline [66]. hsa-miR-223-3p controls both inflammation and ferroptosis regulation by targeting ferroportin genes.Moreover, it’s dysregulation has been linked to increased oxidative stress and neuronal injury in Alzheimer’s and other neurodegenerative diseases [67, 68]. hsa-miR-106b-5p regulates the expression of the APP (Amyloid Beta Precursor Protein) gene. The processing of APP results in the production of amyloid-beta (Aβ) peptides, which build up to form the plaques that are characteristic of Alzheimer’s disease. It also regulates genes related to the cell cycle and has been shown to have neuroprotective effects by preventing the expression of pro-apoptotic factors. A lack of regulation of miR-106b-5p has been linked to cognitive decline in Alzheimer’s disease because of its effects on synaptic plasticity and neurogenesis [69]. mir-125b-5p is the least studied miRNA in context of AD but some structurally and functionally similar miRNAs are found to be associated with genes responsible for regulating tau aggregation and neuroinflammation [70].

Hence, numerous studies have demonstrated the extensive association between miRNAs (hsa-mir-146a-5p, hsa-mir-27a-3p, hsa-mir-106b-5p) and AD, which in turn can be used as a reliable indicator or biomarker for diagnosing AD [61]. Many studies have shown variations of these miRNAs in the plasma of AD patients. This also highlights their significance in coagulation, oxidative stress, and inflammatory pathways, namely tau and neurofilament phosphorylation [71]. Although hsa-mir-125b-5p may include target genes involved in synaptic activity, cell viability and neuroinflammation, but its exact role in AD needs to be further investigated. The gene-miRNA interaction network gives an overall picture of the potential miRNAs which could be used to control the expression of the relevant candidate genes i.e., CYBB and ACSL4 (SI. 4).

### 4. Interplay between biomarkers and immune activation revealed by immune microenvironment analysis

In the previous section we have identified candidate biomarker genes which can be potentially used as therapeutic agents. In this section, we will analyze the immune microenvironment and will identify all the immune cells which are involved in the immune activation process in AD. The heatmap shown in Fig. 7(A) depicts the Pearson correlation coefficients between two AD-related genes i.e., CYBB and ACSL4 and a set of seven selected miRNAs i.e., hsa-miR-146-5p, hsa-miR-106b-5p, hsa-miR-223-3p, hsa-miR-155-5p, hsa-miR-34a-5p, hsa-miR-125b-5p, hsa-miR-27a-3p. A strong negative correlation is observed between CYBB and hsa-miR-106b-5p (r = −0.61) indicating a potential interaction between post-transcriptional regulatory mechanisms. On the other hand, CYBB shows moderate positive correlations with hsa-miR-223-3p (r = 0.27) and hsa-miR-155-5p (r = 0.24). ACSL4 displayed a significant positive correlation only with hsa-miR-106b-5p (r = 0.35) but relatively weak correlation profile across the miRNAs. The gene-gene correlation between CYBB and ACSL4 was highly negative (r = −0.65), indicating opposite expression patterns. This could reflect dissimilar functions or involvement in opposing pathways. Furthermore, inter-miRNA correlations indicated coordinated expression of some miRNAs such as hsa-miR-125b-5p and hsa-miR-34a-5p (r = 0.35), indicating the presence of co-regulatory miRNA modules that may contribute to gene network control in the disease condition. The Fig. 7(B) compares the correlation structure of immune cell populations between healthy controls and AD samples. The heatmaps (left and right) shown in Fig. 7b visualizes the Spearman correlation between various immune cell types in AD compared to healthy controls, highlighting notable differences in immune interactions. The left heatmap represents the raw expression profiles of selected immune cells across all samples emphasizing more on expression intensity and grouping patterns. Rows represent immune cells and columns represent samples. On the other hand, the right heatmap displays the z-score normalized expression matrix, allowing better comparison across immune cells with different expression scales. This standardization helped in highlighting the relative upregulation or downregulation patterns within each sample group and improved visibility of co-expression trends. While both heatmaps contain the same immune cells, the key difference lies in the scaling i.e., the raw heatmap preserved absolute expression levels while z-score heatmap allowed easier pattern detection and immune cell-cell relationship analysis across samples. This dual presentation enables complementary interpretation, one for magnitude and the other for relative changes as well as ensures representation of all immune cell expression in its absolute biological context and its relative regulatory trends and detection of subgroup-specific expression shifts across the AD risk factors. Hence, from both the heatmaps (Fig. 7(B)), we observed a noticeable and organized correlation pattern among immune cell subsets in AD particularly involving macrophages (M0, M1, M2), monocytes and multiple T cell subsets. These patterns suggest a highly dynamic and interactive immune environment which is comparable to the inflammatory response characteristic of AD pathology. Moreover, macrophage populations exhibited strong correlations with both adaptive and innate immune cells underscoring their central role in neuroinflammation. In contrast, healthy control samples displayed a more heterogeneous and weaker correlation profile, indicative of a balanced and homeostatic immune state. These differences reveals significant immune remodeling in AD and support the hypothesis of immune system involvement in disease progression, particularly through altered cellular interactions and activation states.

**Fig. 7.**
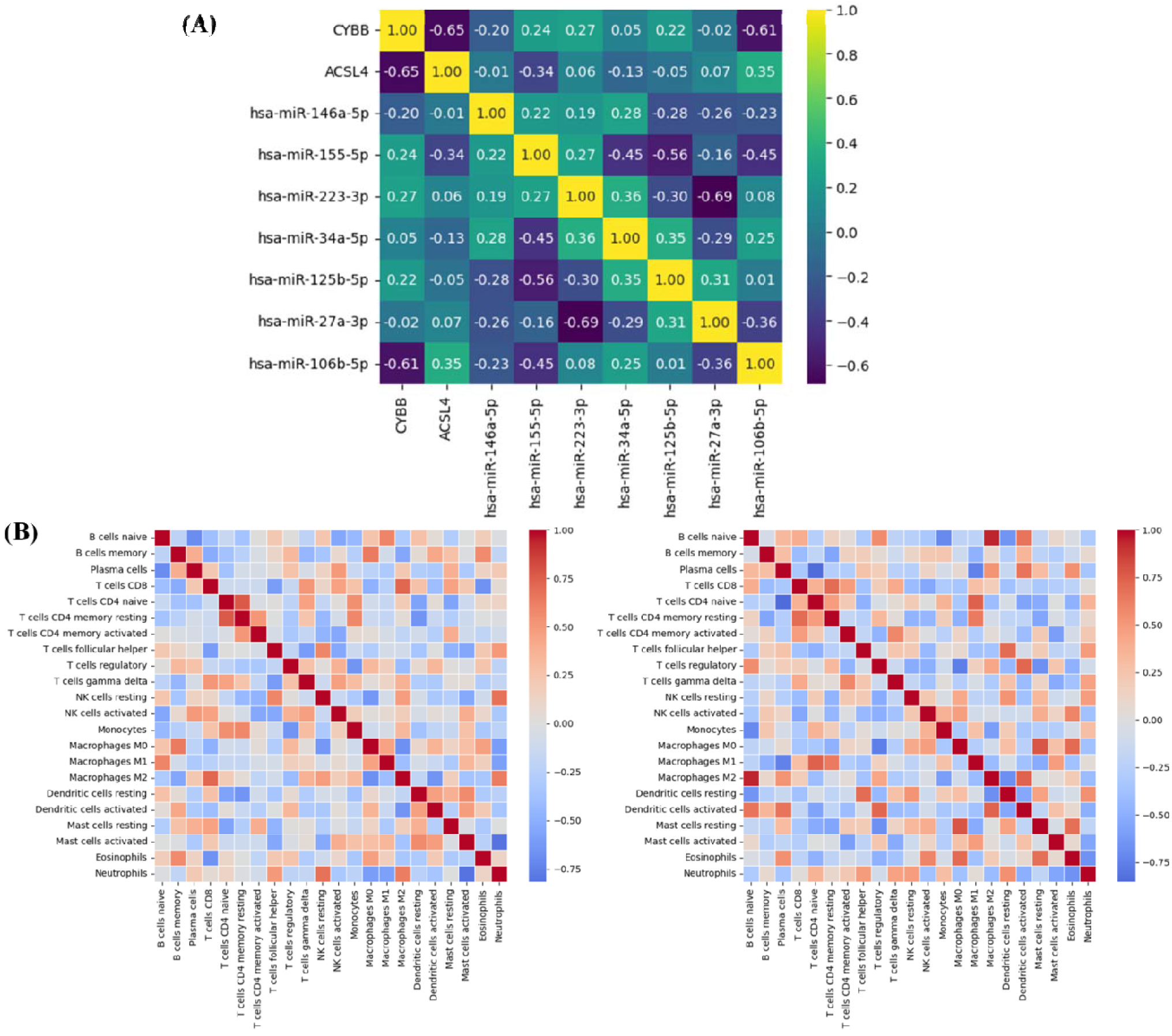
**(A)** Correlation between 22 immune cells, **(B)** Correlation between 22 immune cells and candidate genes.

### 4. Assessing druggability of the identified key biomarker genes and miRNAs in AD pathogenesis

In the previous section some key biomarkers (CYBB and ACSL4) and their regulatory miRNAs were identified. Here, we aim to evaluate the druggability of these identified targets and explore existing drugs that could potentially be repurposed for therapeutic intervention. Using Enrichr-DSigDB, we identified existing drugs which can be used against concerned biomarkers [72, 73]. The top 10 drugs sorted on the basis of decreasing ‘Combined Score’ are shown in Fig. 8.

**Fig. 8.**
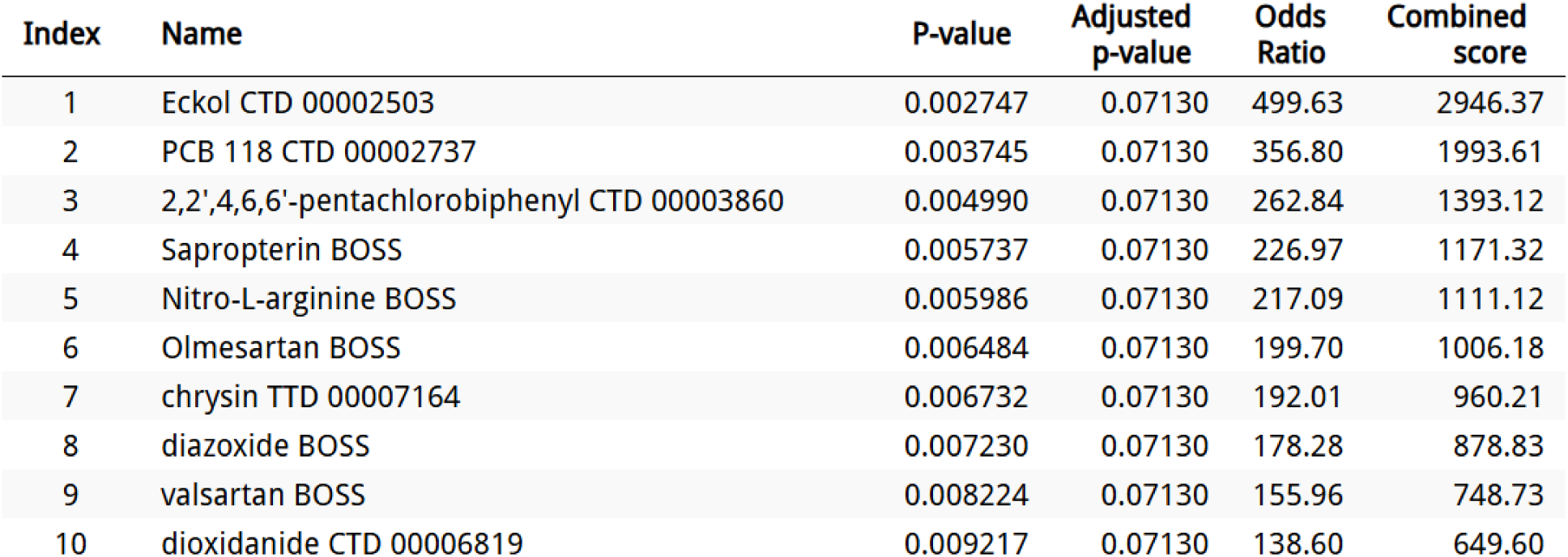
List of top 10 potential repurposed drugs identified for therapeutic intervention in AD pathogenesis

This list provides existing drugs from different sources (like CTD: Comparative Toxicogenomics Database, BOSS, TTD: Therapeutic Target Database) which could be potentially repurposed for target condition and pathway which is AD and ferroptosis. These drugs can help via modulating oxidative stress, iron homeostasis, neuroinflammation or signaling pathways. Here, p value gives a statistical significance of the association between the compound and the target condition i.e., AD and target condition which is ferroptosis. Smaller the value greater is the degree of association. Similarly Odds Ratio also provides the association between drug and target condition. Higher odds ratios are suggestive of stronger associations. Combined scores generally combines all the relevant factors like statistical significance, odds ratio and pathway relevance to rank the compounds according to their priority [73]. ‘Eckol’ is having the highest combined score (2946.37) and a very high odds ratio (499.63), hence ranked first in the list suggestive of its strong potential relevance against AD and ferroptosis. Eckol is a phlorotannin obtained from brown algae which is having antioxidant properties that may protect neuronal cells from oxidative stress which is considered to be the major contributors in ferroptosis and hence AD progression [74]. PCB derivatives are explored to identify their potential to influence signaling and iron-related oxidative pathways under controlled conditions. Similarly, ‘Sapropterin’, a synthetic form of tetrahydrobiopterin (BH4), is essential for neurotransmitter synthesis. Although direct studies on its role in AD are limited, BH4’s role in neurotransmission indicates its potential relevance [75]. ‘Olmesartan’ is another potential drug which acts as an angiotensin II receptor blocker and thus can improves cognitive function and also reduce amyloid-beta accumulation in AD animal models [76].

Furthermore, top compounds are grouped specifically by their primary mechanism of action, focusing on three key ferroptosis-modulating processes: iron chelation and suppression of oxidative stress sand lipid peroxidation as shown in Table 1. These mechanisms directly relate to the functional roles of the identified biomarkers CYBB and ACSL4 in AD pathogenesis.

**Table 1.** Mechanistic clustering of top repurposed drugs.

| **Drug** | **Mechanism of Action** | **Relevance to Ferroptosis** |
| --- | --- | --- |
| **Eckol** | Inhibits NADPH oxidase → ↓ ROS | Decreases the CYBB-mediated oxidative stress |
| **Olmesartan** | Improves iron transport & reduces neuroinflammation | Iron accumulation decreases which decreases ferroptosis |
| **Sapropterin** | Enhances NO signaling and reduces ROS | Supports antioxidant balance, indirectly protects neurons |
| **PCB Derivatives** | Modulates iron storage and peroxidation | Impacts lipid peroxidation which may reduce ACSL4 activation |

## Discussion

This study establishes a mechanistic link between ferroptosis and early-stage AD by integrating transcriptomic data, ML-NN, miRNA networks and immune microenvironment profiling. Enrichment of ferroptosis among DEGs and associated biological processes including mitochondrial function and iron metabolism supports the hypothesis that ferroptotic stress contributes to neurodegeneration in AD. The identification of CYBB and ACSL4 as significantly genes, both functionally linked to oxidative stress and lipid metabolism, underscores their potential as early diagnostic biomarkers. Their regulation by shared hub-miRNAs such as *miR-34a-5p and miR-155-5p* alongwith individual miRNAs such as *miR-146a-5p, hsa-miR-125b-5p and hsa-miR-27a-3p (for CYBB), hsa-miR-106b-5p and hsa-miR-223-3p (for ACSL4),* suggests a tight post-transcriptional control network that could be exploited for therapeutic intervention. These miRNAs are already recognized for roles in neuroinflammation and synaptic dysfunction in AD.

Immune profiling revealed increased infiltration of *macrophages and T cells*, with altered co-expression networks in AD brains. These immune alterations, potentially driven by ferroptosis-induced oxidative damage, align with known inflammatory pathology in AD and could be used to monitor disease progression or therapeutic response. Notably, the druggability assessment identified *Eckol* and *Olmesartan* as promising repurposed agents. Their antioxidant and neuroprotective properties, particularly through inhibition of NADPH oxidase and modulation of lipid peroxidation, present viable routes for targeting ferroptosis in AD.

### Clinical Implications for diagnosis and therapy

The findings from this study offer critical translational value particularly in the early diagnosis and therapeutic targeting of AD. The identification of CYBB and ACSL4 as ferroptosis-linked biomarkers regulated by miRNAs such as hsa-miR-146-5p, hsa-miR-106b-5p, hsa-miR-223-3p, hsa-miR-155-5p, hsa-miR-34a-5p, hsa-miR-125b-5p and hsa-miR-27a-3p opens the possibility for the development of minimally invasive and cost-effective screening or diagnostic tools. These biomarkers can potentially be detected in peripheral fluids such as blood or cerebrospinal fluid (CSF) using techniques like qRT-PCR [77], droplet digital PCR [78] or miRNA profiling platforms [79]. This could support the establishment of biofluid-based screening tools for at-risk individuals enabling early diagnosis before significant neuronal damage occurs. Additionally, the strong diagnostic performance of the neural network-based predictive model (AUC = 0.92, CI = 0.95) suggests that it can be integrated into digital decision-support systems. Such platforms may combine clinical data, gene expression profiles and cognitive assessment results to guide neurologists in early-stage AD identification, particularly in resource-constrained settings where advanced neuroimaging is less accessible and therby improving patient outcomes. In addition to this, the druggability assessment revealed several repurposable drugs including Eckol and Olmesartan, which are known to target pathways involved in oxidative stress, neuroinflammation, iron homeostasis and lipid peroxidation. These drugs may represent viable combinatorial therapies to delay the progression of neurodegeneration in AD patients.

Together, this multi-layered approach establishes the relevance of ferroptosis in AD pathogenesis and also provides a framework for future clinical translation. By combining ML-NN-based prediction, gene–miRNA networks, immune landscape and drugability analysis, this study supports the dual role of identified biomarkers in both diagnostics and therapy highlighting their potential utility in precision medicine frameworks for development of personalized intervention strategies corresponding to individual’s molecular profile.

## Conclusion

AD or Alzheimer’s disease is a widely known neurodegenerative disease that is caused due to damage to the nerve cells. This damage is usually caused due to aggregation of beta-amyloid or tau proteins. It results primarily in the generation of dementia, short-term memory loss and related ailments. Apart from the protein aggregates that are formed, certain molecular changes have also been observed for the AD. This includes synaptic toxicity, autophagy, neuroinflammation, excess generation of reactive oxygen species (ROS), neuronal death and oxidative stress due to an imbalance in metal homeostasis. Metal homeostasis especially that of iron has been deemed very crucial in recent years. Iron-mediated cell death or Ferroptosis is being increasingly observed in several diseases including AD. Accumulation of lipid peroxides, autophagy and metabolism of iron, lipids and aminoacids are few of the indicators of Ferroptosis.

Cognitive assessment, neuropsychological testing, neuroimaging and biofluid assessment are the diagnostic methods that are used for the detection of AD. However, the results obtained are subjective and inconsistent in several instances. Moreover, Neuroimaging techniques lack sensitivity to early stage AD diagnosis and in several instances the detection might involve invasive methodologies such as biopsy. Machine learning has shown very good accuracy in distinguishing between patients with dementia and healthy controls, as well as between various forms of dementia. However, the studies so far have limited data collection and data analysis methods which gives inconsistent diagnosis.

Here, we have used ML and NN to find a link between AD and Ferroptosis. We have taken a universal AD dataset and identified up-regulated genes in AD. The genes were primarily associated with the Ferroptosis pathways. Functionally the genes were linked to oxidoreductase activity, heme binding and iron ion binding. Subsequently, we found a subset of genes that are found in both AD and Ferroptosis. Among these genes, two, namely CYBB and ACSL4, which are involved in generation of reactive oxygen species that contributes to oxidative stress and production of polyunsaturated fatty acids that is linked to lipid metabolism respectively, were found. We found similar results in both supervised and unsupervised subgroup analysis models. In the end, we have identified seven regulatory miRNAs which are found to have a direct association with AD and Ferroptosis. Out of seven, two are shared hub-miRNAs namely *miR-34a-5p and miR-155-5p* and the rest are individual miRNAs such as *miR-146a-5p, hsa-miR-125b-5p and hsa-miR-27a-3p (for CYBB), hsa-miR-106b-5p and hsa-miR-223-3p (for ACSL .*Thus, the AD pathogenesis was significantly predicted by the designed diagnostic model based on the identified genes. Furthermore, we have identified several existing drugs against identified biomarker genes and associated regulatory miRNAs, which could be potentially repurposed to influence the AD pathogenesis. ‘Eckol’ ranked first in the list of drugs with highest p-value, Odds ratio and hence Combined score followed by PCB derivatives, Sapropterin and Olmesartan on the basis on decreasing Combined Scores.

The diagnostic value of CYBB and ACSL4 (involved in ROS production and lipid peroxidation) lies in their role as biomarkers of oxidative stress and Ferroptosis. miRNAs regulating these genes associated with both AD and Ferroptosis, provides an additional layers of molecular information, enhancing early detection when detected in biofluids and diagnostic accuracy for AD when integrated with clinical evaluations. Together, they can form the basis for a comprehensive diagnostic panel for early-stage Alzheimer’s disease detection and personalized therapeutic interventions. Although this study establishes a direct link between Alzheimer’s disease (AD) and ferroptosis, it is important to note that environmental and lifestyle-related risk factors also play a significant role in contributing to the early onset of AD. The future goal of this study is to integrate the impact of all relevant risk factors including genetic, environmental, and lifestyle-related factors, to enhance the accuracy of early-stage Alzheimer’s disease (AD) detection.

## Data Availability

Alzheimer’s disease dataset (GSE118553) is available in Gene Omnibus (GEO) database. Ferroptosis related genes are extracted from FerrDB and MSigDb. We confirm that the data supporting the findings of this study are available within the article.

## Acknowledgements

Authors thank the National Institute of Technology, Warangal, for providing computational facility and MHRD for the research scholar scholarship. Authors thank Prof. Asim Bikas Das, NITW for their helpful comments and suggestions that significantly improved the quality of this work.

## Authors Information

### Contributions

PS: Conceptualization, Problem Modeling, Methodology, Writing and revising SLR: Supervision, Conceptualization, Problem Modeling, Methodology, Writing and revising.

## Ethics Declaration

### Ethical Approval

The authors declare that they are not associated with or involved in any organization or entity that has a financial or non-financial interest in the topics or materials covered in it.

### Consent for Publication

Not Applicable

## Abbreviations

AD: Alzheimer’s disease
ROS: Reactive Oxygen Species
NN: Neural Network
NPS: Neuropsychiatric Symptoms
ML: Machine Learning
AI: Artificial Intelligence
GEO: Gene Expression Omnibus
KEGG: Kyoto Encyclopedia of Genes and Genomes
GSEA: Gene Set Enrichment Analysis
DEG: Differentially Expressed Genes
LIMMA: linear models for microarray data
FDR: False Discovery Rate
LASSO: Least Absolute Shrinkage and Selection Operator
RF: Random Forest
ROC: Receiver Operating Characteristic
AUC: Area Under the Curve
IOBR: Immuno-oncology biological research
AsymAD: Asymptomatic AD
GO: Gene Ontology
SVM: Support Vector Machine
CYBB: Cytochrome b-245 beta chain
AKR1C1: Aldo–Keto Reductase 1
ACSL4: Acyl-CoA synthetase long-chain family member 4
GPX4: Glutathione Peroxidase 4
FTH1: Ferritin Heavy Chain 1
MAPT: Microtubule Associated Protein Tau
CC: Consensus Clustering
CTD: Comparative Toxicogenomics Database
TTD: Therapeutic Target Database
BH4: Tetrahydrobiopterin

