## Supplementary Information for "Neural Network-Enhanced Investigation of Ferroptosis and Druggability in Early-Onset Alzheimer’s Disease"

**SI. 1**

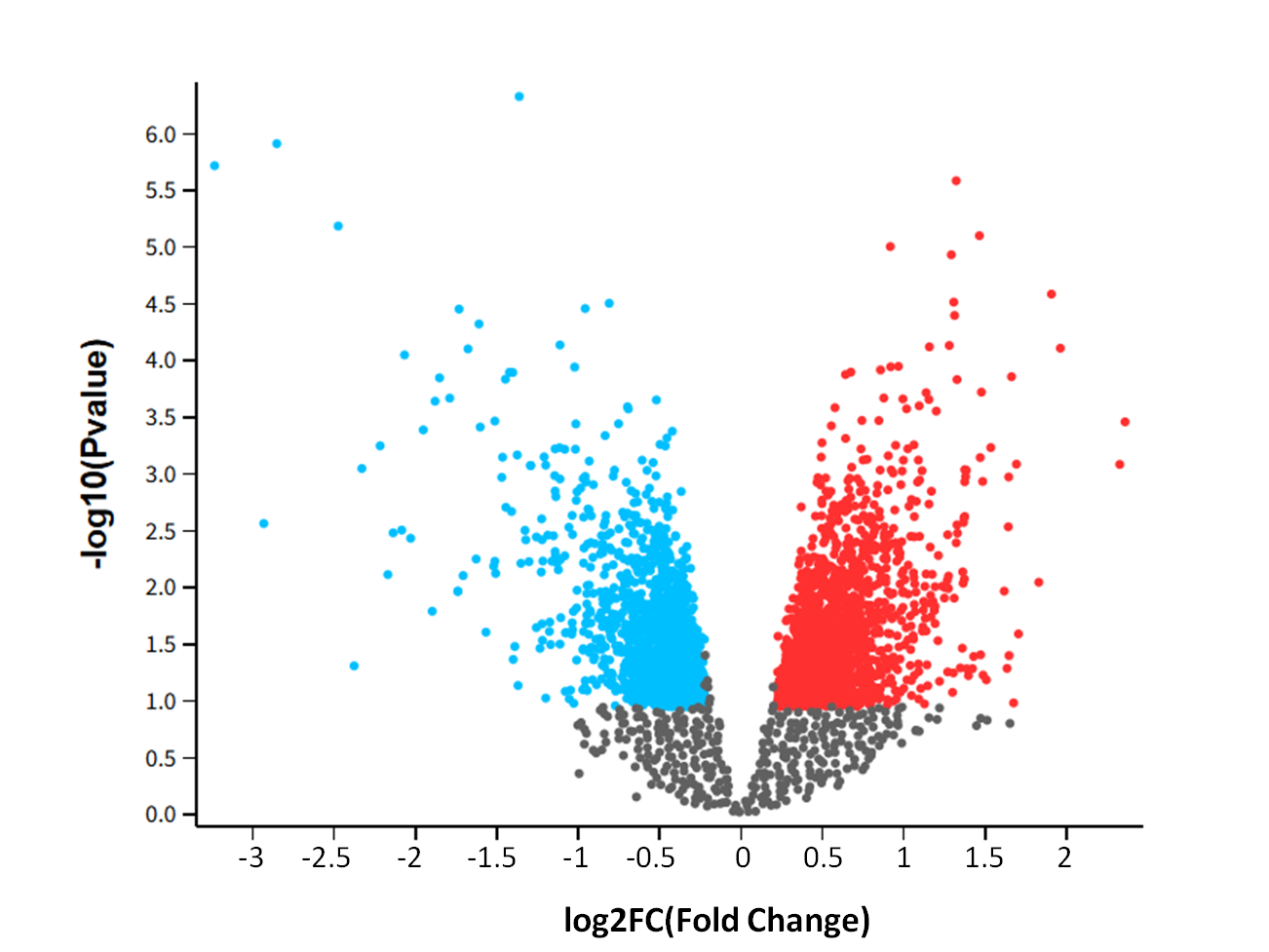

**SI. 1** Volcano map of DEGs in Alzheimer’s disease dataset (GSE118553) where upregulated genes (2107 genes) are shown in red color whereas downregulated genes (1643) are shown in blue color

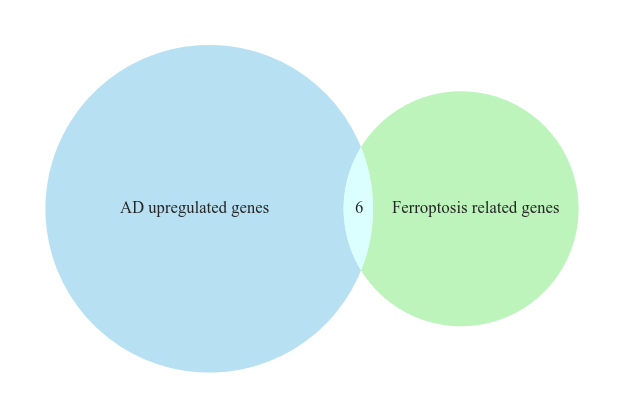

**SI. 2**

**SI. 2** Venn diagram showing crossover between AD differentially expressed genes (47323) obtained from Z-score normalization followed by False Discovery Rate (FDR) correction and Ferroptosis related genes (106). Six genes are common in both the sets and is represented as the intersection between them.

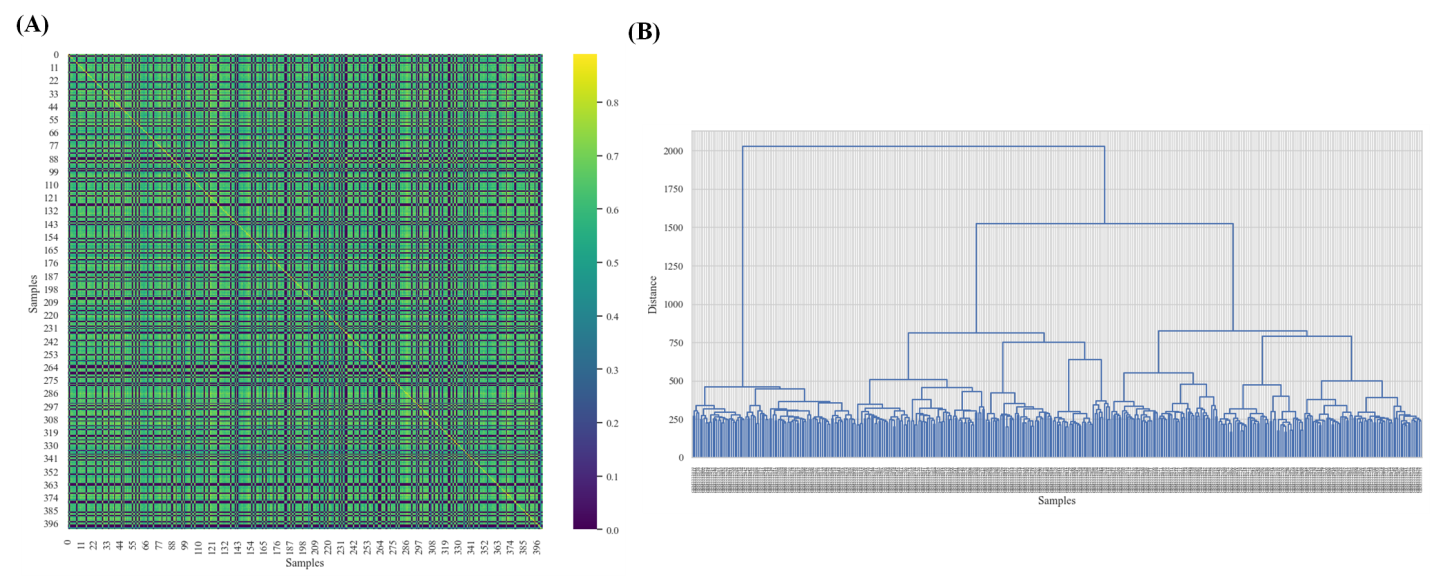
**SI. 3 (A), (B)** Consensus clustering analysis of the relevant genes showing K=2 shown in the form of histogram and dendrogram respectively.

**SI. 3**

**SI. 4**

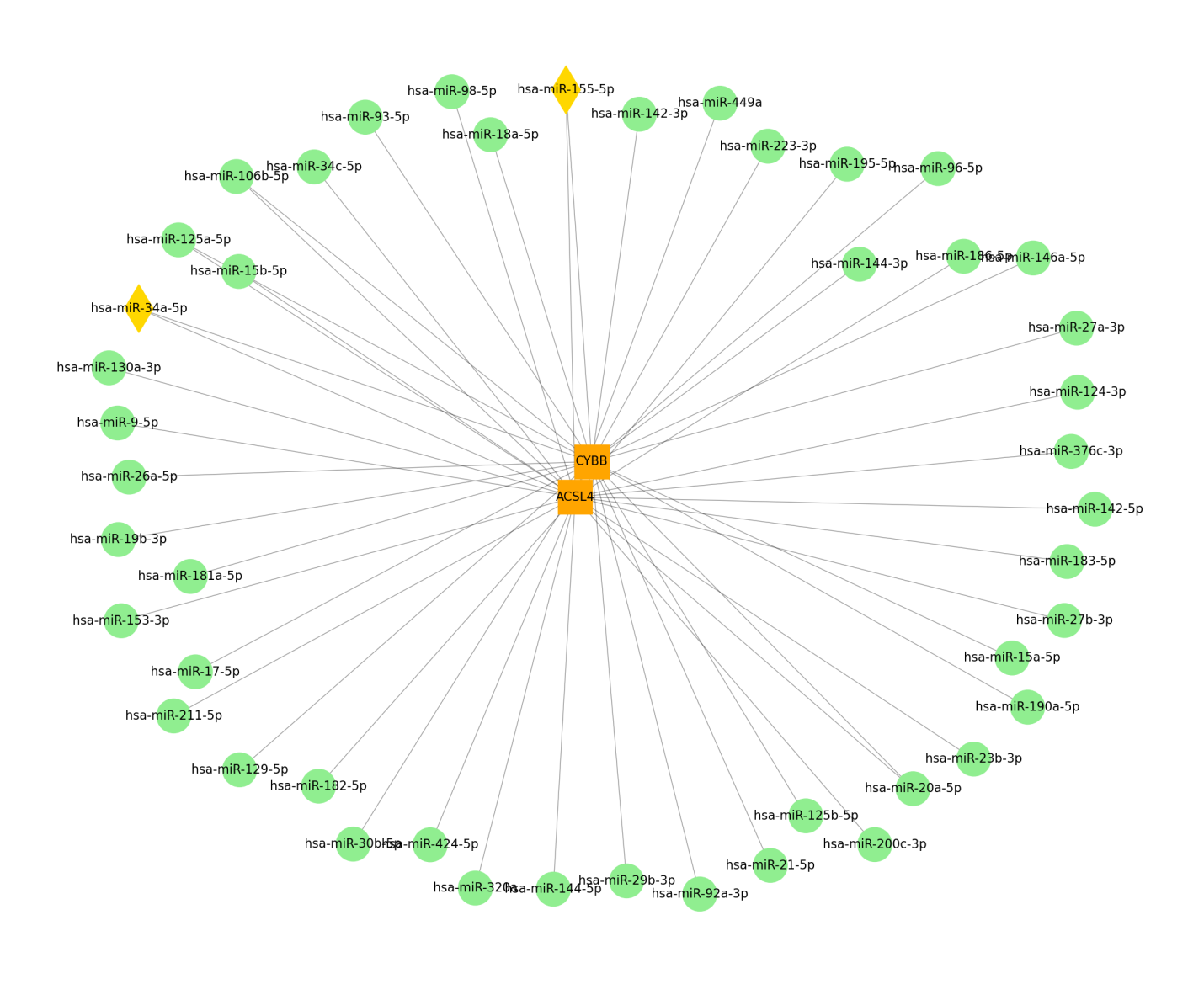

**SI. 4** Elaborated form of Gene-miRNA interaction network showing CYBB and ACSL4 genes in orange color and their regulatory miRNAs in green color alongwith shared miRNAs shown in yellow color.
